# Husbandry and Maintenance of *Carausius morosus* Laboratory Populations

**DOI:** 10.64898/2026.02.19.706905

**Authors:** Macy Ingersoll, Petra Kovacikova, Yousuf Hashmi, Cassandra G. Extavour

## Abstract

*Carausius morosus*, the Indian stick insect, is a slender twig-like insect endemic to India. Though widely introduced through captivity around the world and commonly used in laboratories or kept as a household pet, standardized animal husbandry laboratory protocols are lacking. Here we report detailed laboratory culture conditions for *C. morosus*. We maintain stocks at 23 °C, 70% relative humidity, and a 12:12 hour light-dark photoperiod. This culture has been successfully sustained under these conditions for over two years, with standardized protocols in place for dietary and cage setup conditions. We also report methods for egg and hatchling care to support ongoing experiments with *C. morosus*. These standardized methods improve reproducibility and accessibility, enabling the broader use of *C. morosus* as a laboratory model system for developmental, behavioral, and physiological studies.

**Summary:** This paper outlines detailed protocols for maintaining a *Carausius morosus* laboratory colony, including key procedures for animal husbandry, egg and hatchling care, and an overview of the species lifespan and biological characteristics.

## Introduction

As interest grows in alternative non-mammalian model systems, insects offer an opportunity to explore a wide range of biological diversity while also providing lower-cost research models, simpler care requirements, and fewer regulatory barriers than mammalian models ^1^. Emerging insect model systems make it possible to investigate unique biological phenomena, including for example parthenogenesis in stick insects like *Carausius morosus* ^2^. While the fruit fly *Drosophila melanogaster* has shaped our understanding of developmental and genetic processes since the early 20th century ^3^, other insect model systems such as *C. morosus* present a unique opportunity not only for developmental biology, but also for investigating motor control, locomotion, and neurohormone-regulated systems ^4^. Additionally, within the field of evolutionary developmental biology, hemimetabolous (directly developing) insects remain underrepresented, with the holometabolous (indirectly developing, or undergoing metamorphosis) *D. melanogaster* serving as the most well-studied model insect. Studying directly developing insect species like *C. morosus*, which undergoes several nymphal instar stages before reaching sexual maturity ^5^, allows us to fill key phylogenetic gaps in evolutionary developmental biology research.

*C. morosus* are easy to rear in captivity, making them well-suited for both laboratory research and teaching environments. Their ability to reproduce parthenogenetically ^6^ eliminates the need for complex breeding setups, reducing maintenance demands and simplifying population management. Although successful rearing benefits from controlled environmental conditions, its husbandry remains straightforward and cost-effective compared to many other laboratory models. *C. morosus* feeds readily on common plant material such as ivy (*Hedera helix*), bramble (*Rubus fruticosus*), or rose leaves (*Rosa* spp.).

Standardized rearing protocols for *C. morosus* are critical for producing consistent and reproducible results in research studies. To our knowledge, no in-depth protocols for rearing *C. morosus* in the laboratory have been published to date. In our laboratory, we have successfully established a comprehensive set of standardized protocols for *C. morosus* laboratory cultures including egg, hatchling, and reproductively mature adult rearing. With these methods, we aim to make this insect more accessible to the broader biological research community.

### Protocol

#### Method 1 - Environmental Rearing Conditions

1. The laboratory culture should be maintained in a climate-controlled room or incubator with environmental settings as follows:
  - 23 ºC
  - Relative humidity 70 %
  - Light-Dark photoperiod of 12:12 hours

#### Method 2 - Establishment and Maintenance of Mesh Enclosures

**NOTE:** Mesh insect enclosures with fabric mesh ventilation on three sides are recommended for housing *C. morosus*. An enclosure measuring 24 × 24 × 36 inches is suitable for maintaining up to 50 adults or, when establishing a new colony, up to 500 hatchlings.

1. Select a mesh enclosure of appropriate size. Ensure the container is clean and free of debris before introducing the animals.
2. For establishing a new colony using mature adults, place 25-50 adult females into a clean 24 × 24 × 36-inch mesh enclosure.
3. For the establishment of a new colony using hatchlings, add no more than 500 hatchlings into a clean 24 × 24 × 36-inch mesh enclosure.
4. **Feeding and Maintenance**:
  - 4.1 - Provide fresh foliage such as ivy, bramble, or rose once per week. Ensure all plant material is pesticide-free.
  - 4.2 - Trim stems to fit vessels (small jars or bottles) filled with tap water to maintain leaf hydration.
  - 4.3 - Place the prepared foliage inside the enclosure, making sure leaves are elevated and accessible to the insects. Carefully close the enclosure after placement.
  - 4.4 - During weekly cage cleanings, remove wilted or dried foliage and any accumulated excrement from the bottom of the enclosure. **NOTE:** Maintaining a clean environment is essential for colony health and well-being.
  - 4.5 - Lightly mist each enclosure daily with 1-2 sprays of tap water from a spray bottle to ensure proper hydration.

### Method 3 - Collection and Maintenance of *C. morosus* Eggs

1. Open an enclosure containing adult females.
2. Using a gloved hand, gently sweep the substrates on the bottom of the cage, which contain both eggs and fecal matter, towards the front opening.
3. Carefully transfer the material using a gloved hand to a 15 cm petri dish for egg retrieval.
4. **Egg Retrieval**
  - 4.1 - Using blunt plastic forceps, carefully pick up each egg and place it into a clean 6 cm petri dish. **NOTE:** Mature eggs are protected by a hardened exochorion and are dark brown, resembling plant seeds (Figure1).
  - 4.2 - Label the dish with the date and time of collection for tracking purposes.
  - 4.3 - A 6cm petri dish is sufficient for storing up to 200 eggs. Keep the lid off during incubation to allow ventilation.
  - 4.4 - Approximately one week before the expected hatching window (74-80 days after egg laying; data not shown), replace the Petri dish cover.
  - 4.5 - Begin daily monitoring of egg containers before the expected hatching date to observe emerging hatchlings. **NOTE:** Under the laboratory conditions described in **Method 1**, approximately 60% of eggs are expected to hatch between 74 and 80 days after being laid (data not shown)

**Figure 1.**
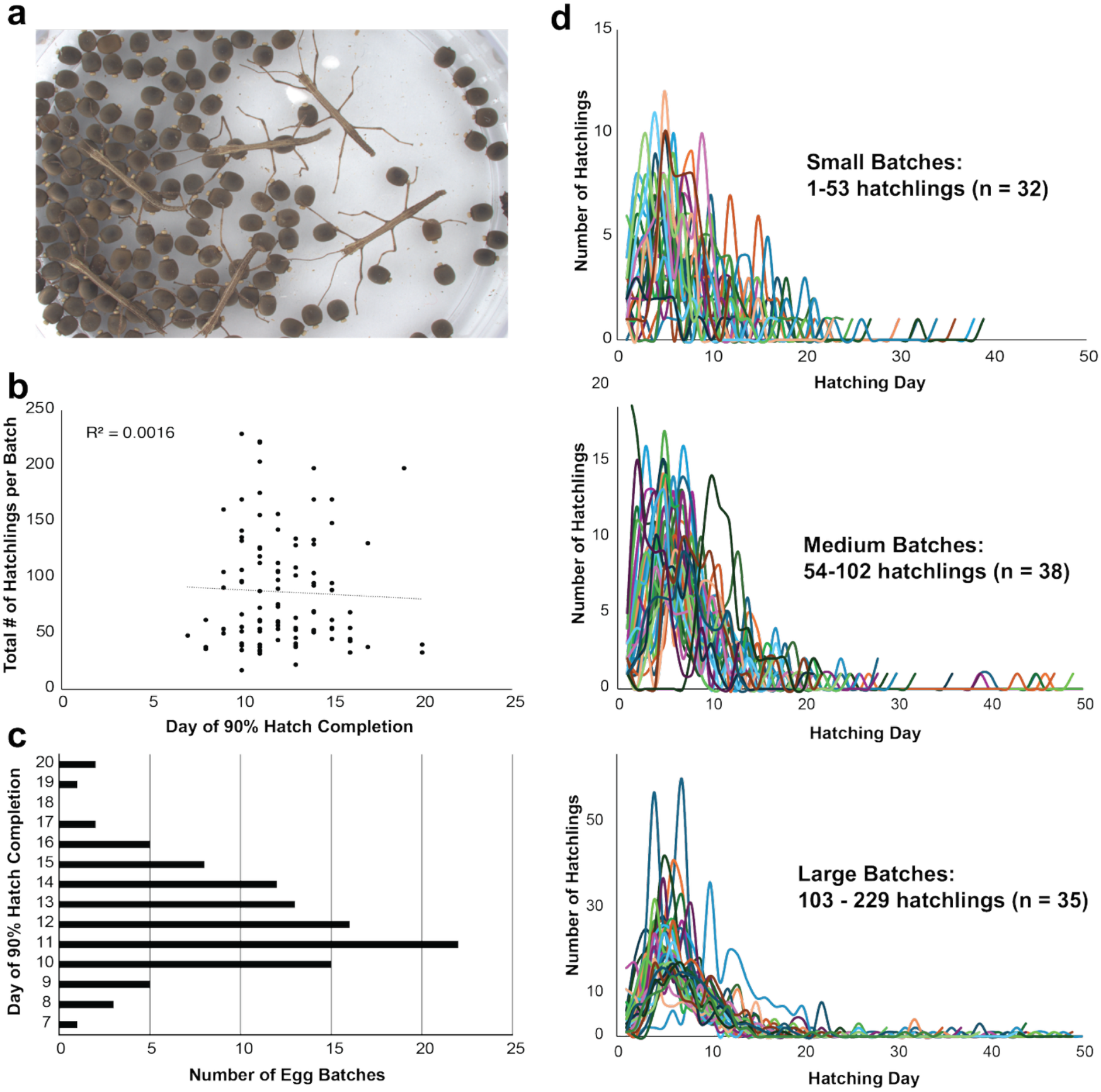
Egg hatching dynamics. (a) Representative image of hatchlings emerging in a Petri dish containing eggs. (b) Relationship between total hatchlings per batch and the number of days required to reach 90% hatch completion. (c). Distribution of egg batches reaching 90% hatch across hatching days 7-20. (d) Hatching trajectories over a 50-day collection period for 105 egg batches collected prior to the establishment of the cages monitored for Figures 2 and 3. Batches were grouped by total hatchling number using percentile cutoffs.

**Figure 2.**
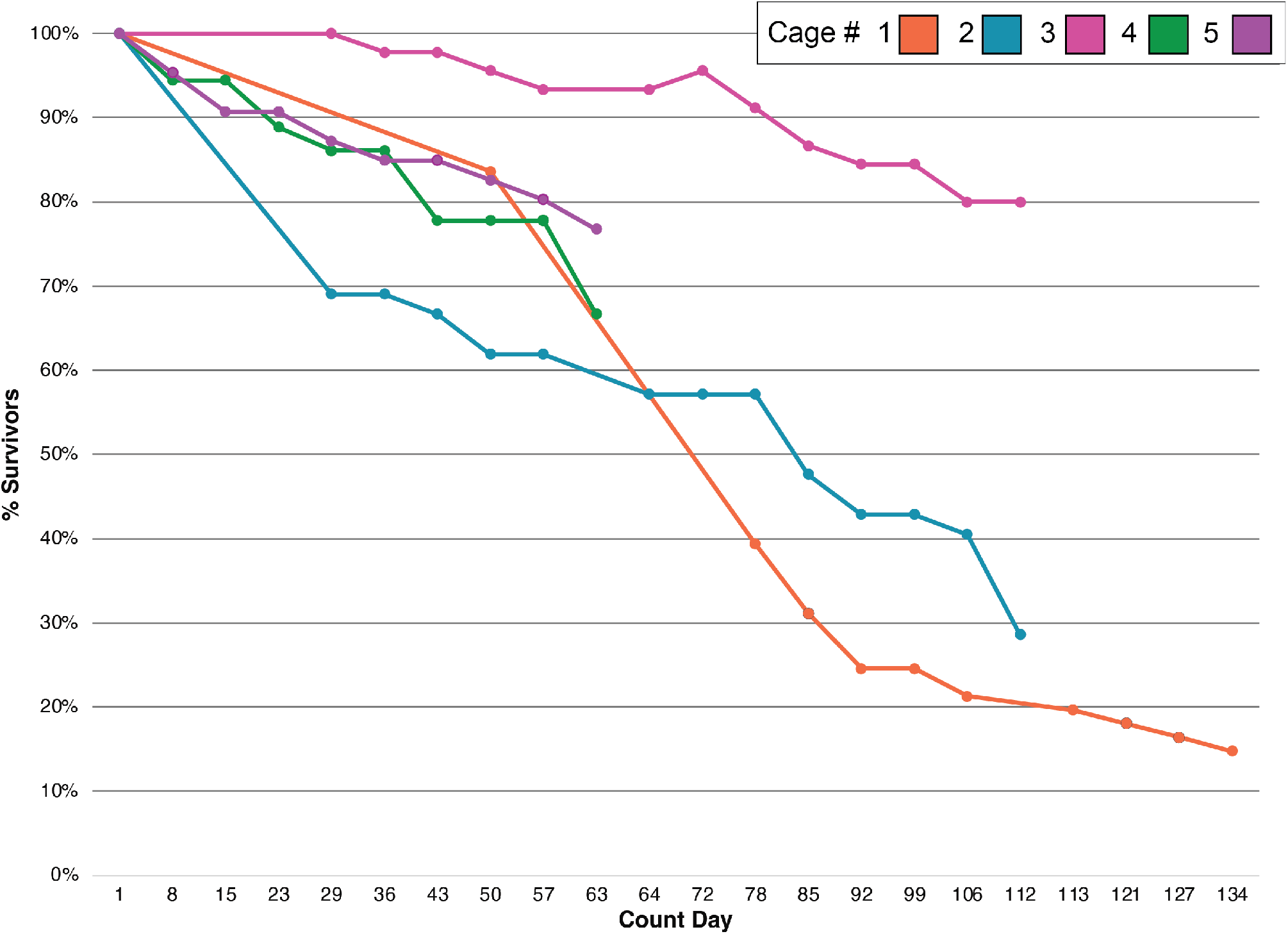
Adult mortality dynamics. Survival trajectories across five replicate *C. morosus* experimental cages over time. Percent survival is plotted over time for individuals in each monitored cage. See summary metrics in Table 1.

**Figure 3.**
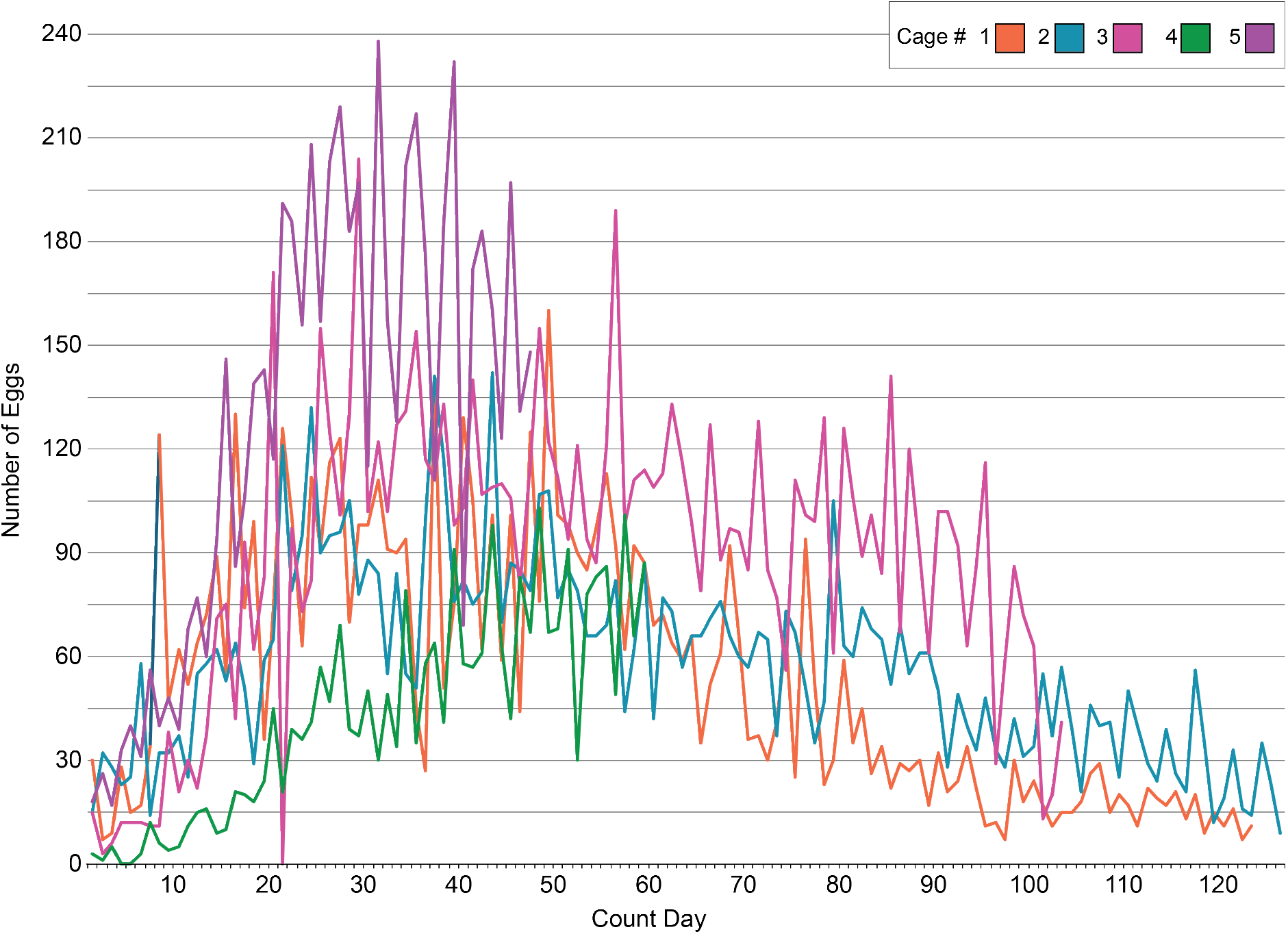
Egg production dynamics. Number of eggs collected from the same five cages monitored in Figure 2. Cages 1 through 5 were monitored daily for the total of 47, 58, 106, 129 and 132 days, respectively. The plot shows data from the first 126 days of oviposition. See summary metrics in Table 2.

### Method 4 - Collection and Maintenance of *Carausius morosus* Hatchlings

1. Gather 6 cm petri dishes with newly emerged hatchlings.
2. Using blunt plastic forceps, gently transfer no more than 500 hatchlings into a clean 24 × 24 × 36-inch mesh enclosure.
3. Provide fresh foliage as described in **Method 2. 4.1 - 4.3**.
4. Label new colony enclosures with the following information:
  - Egg batch collection date(s)
  - Date of hatching or range of hatching dates
  - Number of hatchlings added **NOTE:** This tracks enclosure age and predicts the start of reproductive maturity.
5. Lightly mist each enclosure daily with 1–2 sprays of tap water from an 8oz spray bottle to ensure proper hydration. **NOTE:** Hatchlings may remain in the same enclosure throughout development until adulthood. **NOTE:** Under the laboratory conditions described in **Method 1**, approximately 20% of hatchlings survive to adulthood in 4-5 months (data not shown).
6. Remove any dead or damaged hatchlings during weekly cage cleaning to prevent contamination.

### Method 5 - Life Expectancy

1. Post-hatching, individuals undergo five to six molts before reaching reproductive maturity, which under the laboratory conditions described in **Method 1**, happens around 200 days after hatching (data not shown).
2. Total generation time from egg laying to sexual maturity is approximately 9.5 months (data not shown).

### Method 6 – Termination of Insects and Disposal of Enclosure Waste

When *C. morosus* are no longer needed for experiments, have died, or eggs are not required:

1. Remove insects and any waste materials from the enclosures. Place all contaminated materials into an autoclavable waste receptacle.
  1. Store the waste in a -20ºC freezer for at least 72 hours.
  2. After freezing, transfer the autoclavable waste bag to a biohazard bin for proper disposal.

## Representative Results

We collected data documenting successful laboratory rearing of *C. morosus* from eggs to reproductively mature adults between September 2024 and November 2025. We established a laboratory culture stock during this period from eggs following husbandry steps in **Method 4.1-4.6** of this protocol in a climate-controlled room with a temperature of 23 ºC, 70% relative humidity, and a 12:12 light-dark photoperiod. Over this 14 month period, we established, maintained and monitored five cages for adult mortality, eggs laid, timing of egg hatching, and hatchling survival to adulthood.

Data shown in Figures 2 and 3 present observations from the five monitored cages from setup to adulthood of hatchlings, including the eggs laid and adult mortality for each cage. After the founder individuals in each cage reached sexual maturity and began laying eggs, we recorded the number of adults present in each cage to establish the starting population at the onset of oviposition. Numbers of adults in each cage over time for each cage is shown in **Figure 2** and Table 1. We interpret the steady decline in survival as age-related mortality.

We monitored egg production in parallel in the same five cages monitored for adult survival (Figures 2, 3). Daily egg counts per cage were highest during the first 30-50 days of egg-laying, and decreased gradually over time. Although total egg production varied among cages (Table 2), this general pattern was consistent across all five replicate cages. We monitored egg hatching dynamics over a 50-day period for 105 egg batches (Figure 1d), where each egg batch is a group of eggs oviposited in the same cage within a 24-hour period. We found that once the first egg of a batch had hatched, 90% of eggs from that batch hatched within 12 days of the emergence of the first hatchling (Figure 1d), independent of batch size (Figure 1b). Using the term “hatch completion” to describe the proportion of eggs in a batch that have hatched by a given time point following the emergence of the first hatchling from a batch, we note that the distribution of egg batches by the day of 90% hatch completion indicates that most batches reached this threshold between hatching days 10 and 14 (Figure 1c) after emergence of the first hatchling.

## Discussion

The husbandry methods detailed here offer a straightforward and reproducible framework for maintaining robust laboratory populations of *C. morosus*. Although *C. morosus* has long been kept in laboratories or as domestic pets, laboratory research husbandry protocols have varied across studies ^6, e.g. 7–10^. Ensuring consistent environmental conditions and feeding practices is crucial for making *C. morosus* a reliable laboratory model organism for developmental, physiological, and evolutionary research. Our method supports long-term colony maintenance without signs of disease, crowding-related stress, or reproductive decline. Over more than two years of continuous culture, we observed no obvious pathogen-related mass deaths, and egg and hatchling production remained stable across generations.

### Developmental Timing in C. morosus Relative to Other Model Insects

Compared to holometabolous insects such as the fruit fly *Drosophila melanogaster, C. morosus* exhibits slower developmental rates and a more prolonged juvenile phase. Whereas *D. melanogaster* completes embryogenesis in ∼24 hours and reaches reproductive maturity within 10-12 days at 25°C ^11^, under our rearing conditions *C. morosus* requires 74-80 days for embryogenesis and an additional ∼200 days to reach reproductive maturity at 23ºC (data not shown). This extended timeline reflects fundamental physiological and evolutionary differences between hemimetabolous and holometabolous insects ^12^, and is repeatedly found across the Phasmatodea ^13–15^. While other hemimetabolous insects such as the cricket *Gryllus bimaculatus* complete embryogenesis and reach sexual maturity in a shorter timeframe (ten days and six weeks respectively at 29ºC ^16^), the slower development of *C. morosus* provides a unique opportunity for reproducible research and detailed, longitudinal studies of embryogenesis and juvenile development.

### Colony Size, Survival Rates, and Adult Reproduction

Proper colony setup and careful management of population density were key to keeping the stocks healthy. Maintaining a maximum of approximately 500 hatchlings or 50 adults per 24 × 24 × 36-inch enclosure ensured stable conditions, a healthy colony and reproducible developmental observations.

Comparing average daily mortality with the number of founding adults in a cage (Figure 4a; Table 1) revealed no predictable relationship between these two parameters. Similarly, there was not a strong predictive relationship between total number of founding adults and total egg production (Figure 4b; Tables 1, 2). Instead, cages with lower average daily adult mortality showed a mild tendency to have higher average daily egg output (Figure 4c). However, differences among cages suggest that egg production varied per adult (Tables 1, 2), which could mean that factors beyond adult survival contribute to fecundity patterns, and that individual females may also vary in how many eggs they lay each day.

**Figure 4.**
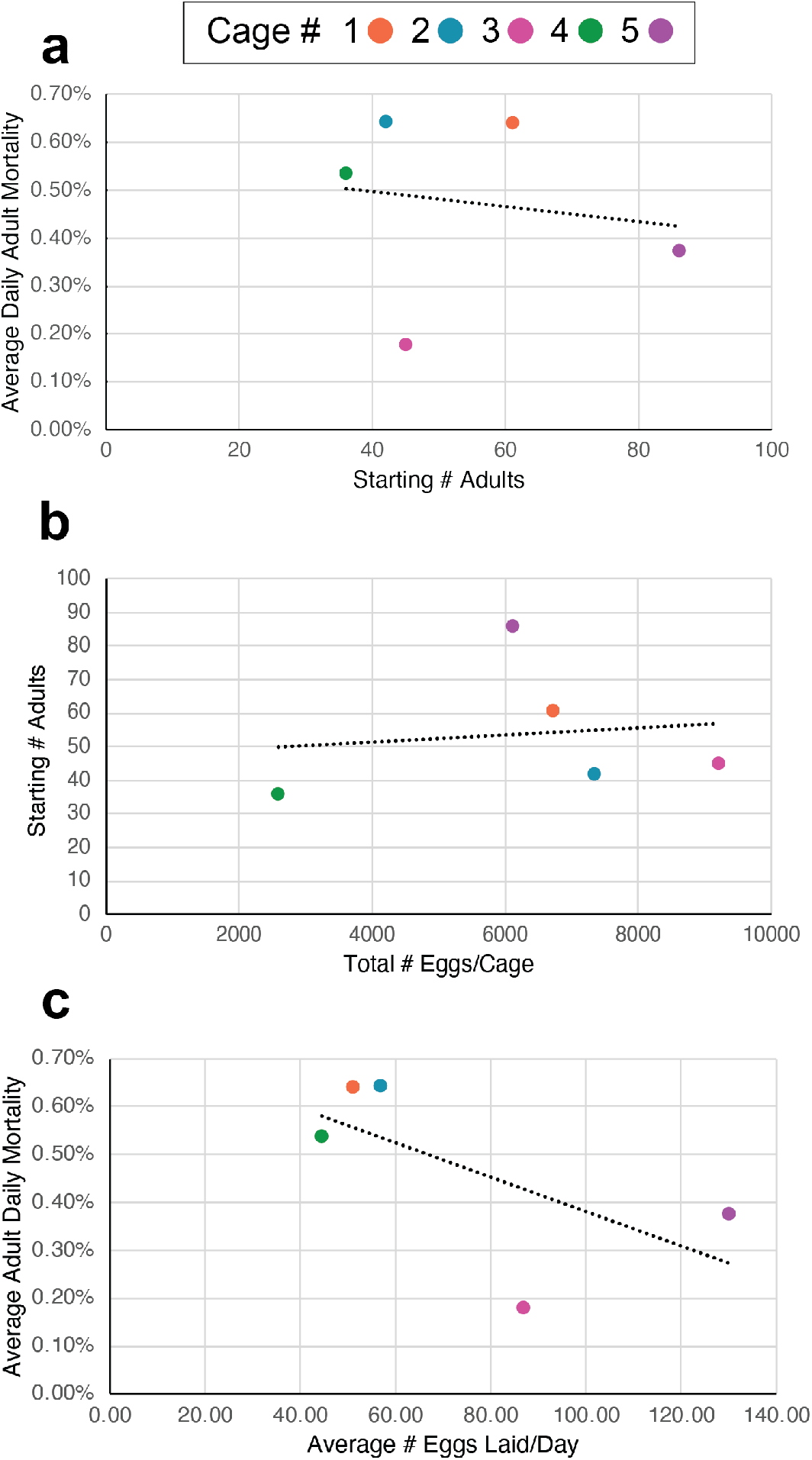
Relationships between adult numbers, adult mortality, and egg production per cage. (a) There is no significant relationship (R^2^ = 0.025) between average adult daily mortality and the number of founding adults in a cage. (b) There is no significant relationship (R^2^ = 0.017) between total egg production and the number of founding adults in a cage. (c) There is a moderate negative correlation (R^2^ = 0.411) between average daily egg production and average adult daily mortality. Average adult daily mortality = (% total mortality) / (total number of days monitored)

Egg production in all monitored cages generally declined over the course of the experiment (Figure 3). Most cages had the highest daily egg-laying rates early in the reproductive period (first 50 days), after which production gradually tapered off (Figure 3). This pattern is consistent with our observation that individuals typically reach sexual maturity and begin laying eggs around 135 days post-hatching, and mortality starts approximately 90 days later (∼225 days post-hatching; data not shown).

### Timing and Dynamics of Hatching

Across 105 monitored egg batches, most hatching occurred within a consistent 12-day period following the onset of emergence (Figure 1c), regardless of the number of eggs in a batch (Figure 1d). Similarly, there was no correlation between batch size and the timing of 90% hatch completion (Figure 1b). This predictable hatching period is a useful feature of *C. morosus* colony management because it allows hatchlings to be grouped into cohorts of similar age, improving developmental staging and reducing experimental variability. Knowing when bulk hatching will occur also allows for better planning of enclosure turnover and resource allocation, including the timely preparation of enclosures, foliage, and tracking materials, an especially important consideration for large-scale or long-term studies.

### Egg Handling and Substrate Considerations

Our egg-handling method involves collecting eggs directly from the enclosure floor and incubating them in ventilated Petri dishes. This approach maintains high viability while making eggs easily observable, unlike cricket systems that collect eggs in substrates such as coconut fiber or cotton wicks ^17^. The hard exochorion of *C. morosus* eggs provides protection during handling and against desiccation, making it possible to incubate eggs safely in Petri dishes without additional substrate, while maintaining high hatch rates. Taken together, these standardized procedures provide a structured approach for maintaining healthy *C. morosus* populations for reproducible developmental, behavioral, genetic, or physiological research. Consistent environmental controls (23 °C, 70% RH, 12:12 LD), regular enclosure maintenance, and predictable developmental timelines make *C. morosus* an accessible and low-cost model system. Importantly, unlike some cricket systems where crowding, cannibalism, or pathogen load can rapidly destabilize colonies ^18^, *C. morosus* are easy to maintain, making them well suited for large-scale experiments and studies spanning multiple generations. As *C. morosus* becomes more widely adopted as a hemimetabolous model species, use of standardized methods should promote reproducibility and comparability across studies in multiple areas of insect biology.

## Supporting information

Video Protocol

## Acknowledgments

This project was supported by National Science Foundation Award IOS 22207477 to CGE, who is an investigator of the Howard Hughes Medical Institute. We sincerely thank Stanislav Gorb and Thies Büscher (University of Kiel) for providing the initial group of adult females to establish our culture, and all members of the Extavour lab for their support, including former lab member Dr. Upendra Bhattarai (current affiliation: Brown University) who set up the initial colony.

## Table of Materials Used in Article

| **Material** | **Company** | **Catalog Number** | **Comments** |
| --- | --- | --- | --- |
| 8oz Spray bottle | Amazon | B000H88PCU | For misting mesh enclosures |
| Autoclavable polypropylene bags, W x H=30.5 x 61 cm (12 x 24) | VWR® | 95042-554 | For disposal |
| Disposable Petri dishes 150 x 15 mm | VWR® | 25384-326 | For egg collection |
| Disposable Petri dishes 60 x 15 mm | VWR® | 25384-092 | For egg incubation |
| Ivy, bramble, rose leaves | N/A | N/A | Collected outside |
| Mesh insect enclosures | Amazon | B09GS2CH1G | Used to house insects |
| Tall glass container | N/A | N/A | Used to propagate and hold plant leaves |
| Tap water | N/A | N/A | For misting |
| Utoolmart 10pcs anti-static tweezers | Amazon | B07YZ5FB27 | Used for collecting and handling eggs and hatchlings |

## Table Legends Table Download Link

**Table 1. Adult survival and mortality dynamics across experimental cages.** Adult survival outcomes for five replicate experimental cages. For each cage, we report the date range of cage establishment, the time between cage establishment and initiation of adult counts, the duration of adult monitoring, starting and ending adult population sizes, adult mortality over the monitored period, and average daily mortality.

**Table 2. Egg production across experimental cages.** Egg-laying output of five replicate experimental cages containing mature adults. For each cage, we report the start and end dates of egg collection, the total duration of egg collection, cumulative egg production per cage, and the average number of eggs laid per day.

**High Resolution Video Protocol Download Link** .[mov](https://www.dropbox.com/scl/fi/tmai7306ikrexigunrnua/Ingersoll_Video_V4_FINAL.mov?rlkey=6htkask3ay5r87li1bdtmxy6k&dl=0)

**High Resolution Video Protocol Download Link** .[mp4](https://www.dropbox.com/scl/fi/87de3g7oxzvcyzmy631wv/Ingersoll_Video_V4_FINAL.mp4?rlkey=ucss4059yzomeui8t65oz13r4&dl=0)

